# Stochastic simulation tools and continuum models for describing two-dimensional collective cell spreading with universal growth functions

**DOI:** 10.1101/052969

**Authors:** Wang Jin, Catherine J. Penington, Scott W. McCue, Matthew J. Simpson

## Abstract

Two-dimensional collective cell migration assays are used to study cancer and tissue repair. These assays involve combined cell migration and cell proliferation processes, both of which are modulated by cell-to-cell crowding. Previous discrete models of collective cell migration assays involve a nearest-neighbour proliferation mechanism where crowding effects are incorporated by aborting potential proliferation events if the randomly chosen target site is occupied. There are two limitations of this traditional approach: (i) it seems unreasonable to abort a potential proliferation event based on the occupancy of a single, randomly chosen target site; and, (ii) the continuum limit description of this mechanism leads to the standard logistic growth function, but some experimental evidence suggests that cells do not always proliferate logistically. Motivated by these observations, we introduce a generalised proliferation mechanism which allows non-nearest neighbour proliferation events to take place over a template of *r* ≥ 1 concentric rings of lattice sites. Further, the decision to abort potential proliferation events is made using a *crowding function, f* (*C*), which accounts for the density of agents within a group of sites rather than dealing with the occupancy of a single randomly chosen site. Analysing the continuum limit description of the stochastic model shows that the standard logistic source term, λ*C*(1 – *C*), where λ is the proliferation rate, is generalised to a universal growth function, λ*Cf* (*C*). Comparing the solution of the continuum description with averaged simulation data indicates that the continuum model performs well for many choices of *f* (*C*) and *r*. For nonlinear *f* (*C*), the quality of the continuum-discrete match increases with *r*.

## I. INTRODUCTION

Two-dimensional collective cell migration assays are routinely used to study combined cell migration and cell proliferation processes [1]. These assays provide insight into cancer [2] and tissue repair [3]. There are two different kinds of cell migration assays: (i) *Cell proliferation assays*, as shown in Figure 1(a)–(d), are initiated by uniformly distributing cells on a two-dimensional substrate. Over time, individual cells undergo migration and proliferation events, leading to the formation of a confluent monolayer [4, 5]; and, (ii) *Scratch assays*, as shown in Figure 1(e)–(h), are initiated in the same way as a cell proliferation assay, except that a wound, or scratch is made in the monolayer [6]. In a scratch assay, individual cells undergo motility and proliferation events with the net result being the spreading of cells into the vacant region [6]. A critical feature of collective cell migration assays is the role of crowding. At low cell density, individual cells are relatively free to move and proliferate because of the abundance of free space [4, 7]. In contrast, at high cell density, individual cells are strongly influenced by cell-to-cell crowding, which reduces their ability to move and proliferate [4, 7].

**Figure 1.**
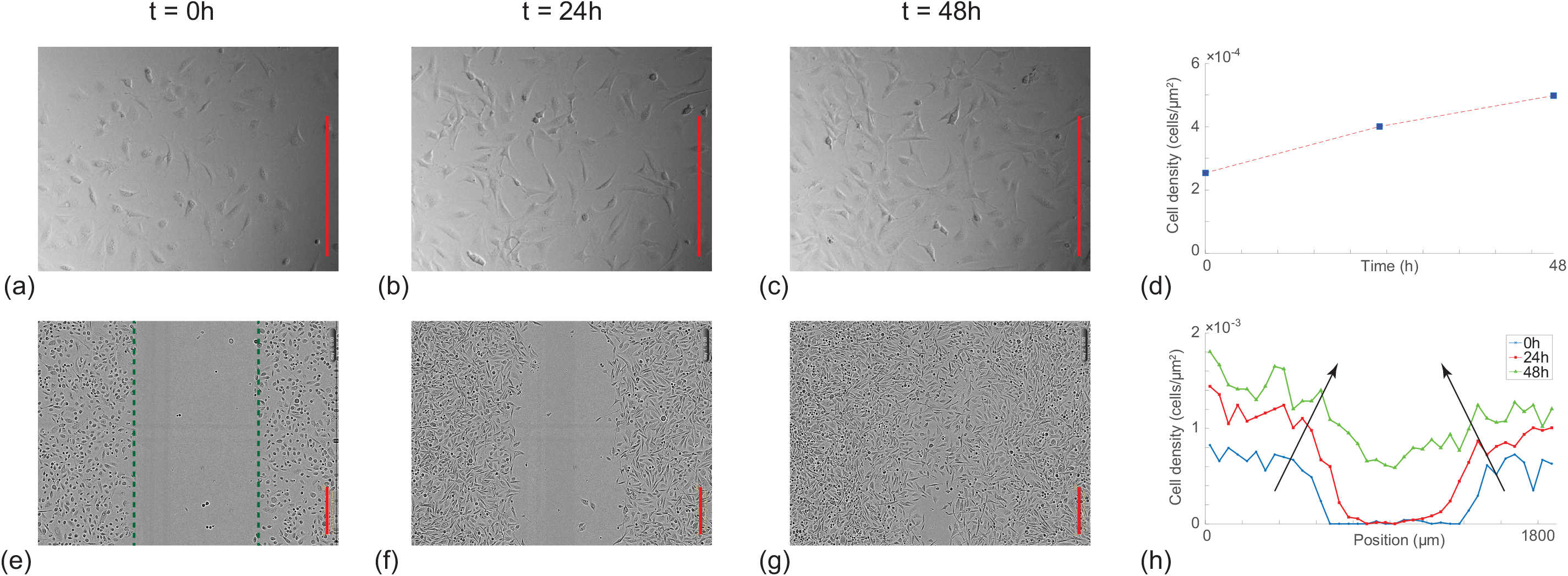
Experimental motivation. (a)-(c) Images from a *cell proliferation assay*, shown at *t* = 0,24 and 48h, respectively [5]. The cell proliferation assay is initiated by uniformly distributing 25,000 3T3 fibroblast cells into the wells of a 24-well tissue culture plate [5]. The dimension of the field of view is 640*μ*mx480*μ*m, and the spatial extent of the growing population extends well beyond the field of view [5]. Results in (d) shows the increase in cell density as a function of time obtained by counting the number of cells in the images in (a)-(c). (e)-(g) Images from a *scratch assay*, shown at *t* = 0, 24 and 48h, respectively [6]. Experiments are initiated by uniformly distributing 16,000 PC-3 cells into the well of a 96-well tissue culture plate [6]. A scratch (dashed green in (e)) is made at *t* = 0 [6], and the subsequent healing of the wound is observed with time. The dimension of the field of view is 1900/*μ*m x 1400*μ*m, and the spatial extent of the population extends well beyond the field of view [6]. The plot in (h) shows the spatial distribution of cell density obtained by discretising the images into strips of width 50 *μ*m and counting the number of cells per strip. Dividing the number of cells in each strip by the area of the strip gives an estimate of the cell density. Three plots are given in (h) showing the cell density profile at *t* = 0, 24 and 48 hours, with the arrows indicating the direction of increasing time. All scale bars correspond to 300*μ*m.

There are two different approaches to modelling collective cell migration assays. Firstly, a continuum reaction-diffusion equation can be applied to mimic certain features of the experiment [3]. Most previous continuum models represent cell migration with a diffusion-type mechanism, and a logistic source term to represent carrying capacity-limited proliferation [3, 7–11]. Secondly, a discrete random walk model can be used to mimic certain features of the experiment [12, 13]. Here, many previous studies represent cell migration using an unbiased exclusion process [14], which incorporates hard core exclusion to model cell-to-cell crowding [15–20]. Cell proliferation is incorporated by allowing agents to place daughter agents on the lattice, with crowding effects incorporated by ensuring that potential proliferation events that would place a daughter agent on an occupied site are aborted [17, 18]. Discrete random walk models have an advantage over continuum models when it comes to comparing model predictions with experimental observations. Experimental data involving individual cells can be directly compared with the predictions of discrete models, whereas continuum models do not provide direct information about individual cells [21, 22].

Lattice-based random walk models of collective cell migration assays typically involve a particular mechanism to assess how crowding influences potential proliferation events [15–18]. Mean field analysis of this traditional proliferation mechanism leads to the logistic source term in the partial differential equation (PDE) description of the model [5, 17, 18]. In terms of continuum models, carrying-capacity limited proliferation is often represented using the logistic equation, d*C* / d*t* = λ *C*(1 – *C*) [3, 7–11], where λ is the proliferation rate, and the density has been scaled relative to the carrying capacity. However, there is some awareness that the logistic model does not always match experimental data. For example, West and colleagues [23] examine data describing *in vivo* growth, showing that their data is best described by a generalised logistic model, d*C*/d*t* = λ *C*^3/4^ (1 – *C*^1/4^). Similarly, Laird [24] examines *in vivo* tumour growth data, showing that the dynamics is better described by a Gompertz growth law than the classic logistic growth model. More recently, our previous analysis of a suite of scratch assays suggests that when a logistic-type reaction-diffusion equation is calibrated to match experimental data with a range of initial cell densities, there is no unique choice of λ for which the logistic model matches the entire data set [6]. One way of interpreting this result is that cells do not proliferate logistically. While several previous theoretical studies have analysed generalised logistic growth models, such as d*C*/d*t* = λ*C^α^*(1 – *C^β^*)^*γ*^ for arbitrary positive constants *α, β* and *γ* [25], it is presently unknown how to implement this generalised proliferation mechanism in an exclusion process.

The aim of this work is to analyse a discrete model of two-dimensional collective cell migration assays. In all cases we consider the cell migration to be modelled as an unbiased nearest neighbour exclusion process where potential migration events occur with probability *P_m_* per time step of duration *τ*. This motility mechanism is able to capture certain features of previous *in vitro* experimental data [5, 26]. The focus of our work is on the details of the proliferation mechanism. Potential proliferation events occur with probability *P_p_* per time step of duration *τ*. In the traditional model, the location of the daughter agent is chosen by randomly selecting a nearest neighbour site. If the randomly selected target site is vacant, the proliferation event is successful, whereas if the randomly selected target site is occupied, the proliferation event is aborted [5, 17, 18]. Two extensions of the standard proliferation model are analysed: (i) we consider non-nearest neighbour proliferation mechanisms, whereby the crowdedness of any individual agent is influenced by a larger template on the lattice, and it is possible for the daughter agent to be placed on a non-nearest neighbour site [27, 28]; and, (ii) we adjust the way that we measure the local density, *Ĉ* ∈ [0,1], and implement a new way of deciding whether to abort potential proliferation events due to crowding by using a more general *crowding function, f* (*C*). We derive the continuum limit PDE description of the generalised discrete model, and apply both the discrete and continuum models to mimic a suite of cell proliferation and scratch assays. Our results illustrate several interesting features about the relationship between the discrete model and the continuum limit description.

## II. DISCRETE MATHEMATICAL MODELS

We adopt the convention that dimensional variables are primed and non-dimensional variables are unprimed. A lattice-based random walk model will be used to describe the collective motion of a population of cells with an average cell diameter of Δ′. The lattice spacing is taken to be equal to the average cell diameter so that there are, at most, one agent per site. All simulations are non-dimensional in the sense that they are performed on a hexagonal lattice with unit lattice spacing, Δ = 1. These non-dimensional simulations can be used to model any particular cell population by re-scaling with the dimensional cell diameter, Δ′. Each lattice site, indexed (*i,j*) where 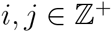, has position

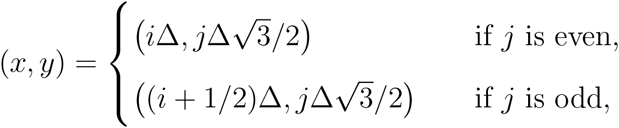

such that 1 ≤ *i* ≤ *I* and 1 ≤ *j* ≤ *J*. In any single realisation of the model, the occupancy of site s is denoted *C*_s_, with *C*_s_ = 1 if the site is occupied, and *C*_s_ = 0 if vacant. Since site s is associated with a unique index (*i,j*), we will use *C*_s_ and *C_i,j_* interchangeably.

*Traditional discrete model*: If there are *N*(*t*) agents at time *t*, then during the next time step of duration *τ*, *N*(*t*) agents are selected independently at random, one at a time with replacement, and given the opportunity to move [5, 18]. The randomly selected agent attempts to move, with probability *P_m_*, to one of the six nearest neighbour sites (Figure 2(a)), with the target site chosen randomly. Motility events are aborted if an agent attempts to move to an occupied site.

**Figure 2.**
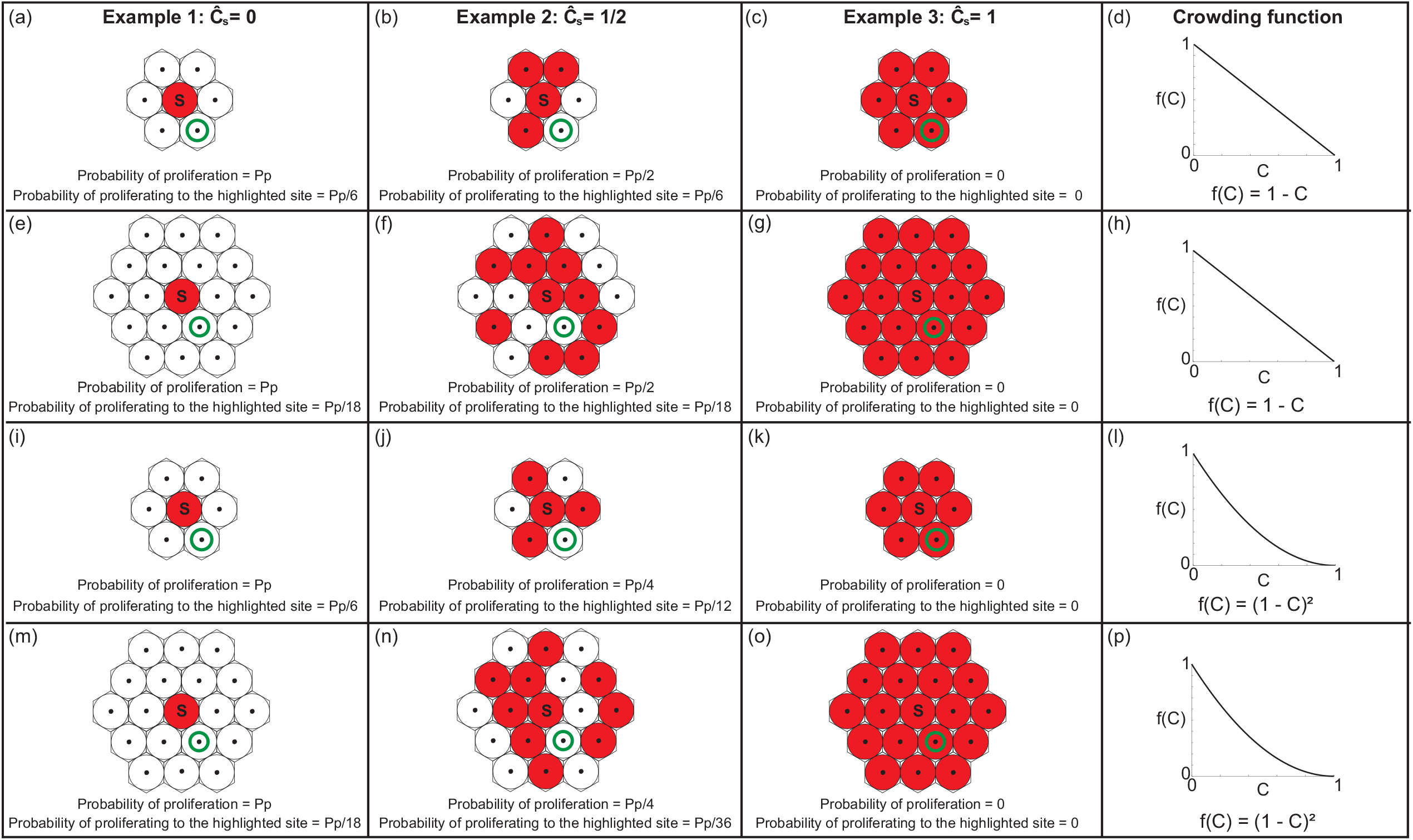
Schematic representation of the proliferation mechanisms considered in this work. Three examples are illustrated, in the first three columns, for *Ĉ* = 0,0.5 and 1, respectively. In each lattice fragment, the central site s is occupied (red), while some of the neighbouring sites are occupied (red) and others are vacant (white). In all cases we always consider the outcomes for a potential proliferation event of the agent at the central lattice site. The properties associated with the traditional proliferation mechanism are illustrated in (a)-(d). The properties associated with the first generalisation, where we consider the same crowding mechanism as the traditional model, but over a larger template of neighbouring lattice sites, is shown in (e)-(h). The properties associated with the second generalisation, where we consider the same nearest neighbour template as the traditional model, but we consider a different method of aborting potential proliferation events using *f*(*C*), is shown in (i)-(l). The final row, (m)-(p), illustrates how the two generalisations can be combined. To highlight differences between the generalisations we report: *Ĉ*_s_; the probability of a successful proliferation event taking place; and, the probability of a proliferation event successfully depositing a daughter at a particular site, highlighted with a green annulus, in (a)-(c), (e)-(g), (i)-(k) and (m)-(o).

Once *N*(*t*) potential motility events are attempted, another *N*(*t*) agents are selected independently, at random, one at a time with replacement, and given the opportunity to proliferate with probability *P_p_*. The location of the daughter agent is chosen, at random, from one of the six nearest neighbour sites [5, 17, 18]. If the selected site is occupied, the potential proliferation event is aborted. In contrast, if the selected site is vacant, a new daughter agent is placed on that site. After the *N*(*t*) potential proliferation events have been attempted, *N*(*t* + *τ*) is updated [5, 17, 18]. One of the limitations of the traditional discrete model is that the continuum limit description leads to the traditional logistic source term [17, 18], however certain experimental observations indicate that logistic growth is not always appropriate [6, 23]. As such, we consider two extensions of the discrete proliferation mechanism.

*Extension 1*: We first generalise the traditional discrete proliferation mechanism so that crowding effects are felt over a larger spatial template, and daughter agents can be placed on non-nearest neighbour sites. The placement of daughter agents on non-nearest neighbour sites is consistent with previous *in vitro* [27] and *in vivo* [28] experimental observations. To achieve this, we consider a proliferative agent at site s, and we use 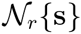 to denote the set of neighbouring sites, where *r* ≥ 1 is the number of concentric rings of sites surrounding s. For example, when 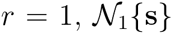 denotes the set of six nearest-neighbouring sites, as demonstrated in Figure 2(a). In contrast, when 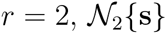 also includes the set of the next nearest-neighbouring sites, as demonstrated in Figure 2(e). More generally, the number of sites in 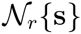 is 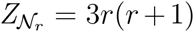. To implement this extension we first choose a value of *r*. For any potential proliferation event, the target site for the placement of the daughter agent is chosen from 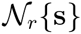. If a randomly chosen target site is vacant, a daughter agent is placed at that site. If a randomly chosen target site is occupied the potential proliferation event is aborted.

*Extension 2*: Instead of deciding to abort a potential proliferation event depending on the occupancy of a single randomly chosen site, we consider a more general approach by assuming that a proliferative agent at site s senses the occupancy of all sites within 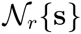, and detects a measure of the average occupancy of those sites, 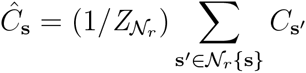. This means that *Ĉ*_s_ ∈ [0,1] is a measure of the crowdedness of the region surrounding s. We anticipate that using C_s_ to determine whether potential proliferation events are aborted is more realistic than the traditional model where the decision depends solely on the occupancy of a single, randomly chosen site. To use *Ĉ*_s_ to determine whether a potential proliferation event succeeds, we introduce a *crowding function, f* (*C*) ∈ [0,1] with *f* (0) = 1 and *f* (1) = 0. To incorporate crowding effects we sample a random number, *R* ~ *U*(0,1). If *R* > *f* (*Ĉ*_s_), a daughter agent is placed at a randomly chosen vacant site in 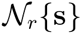, whereas if *R* > *f* (*Ĉ*_s_), the event is aborted. This extension can be applied to different sized templates by varying *r*.

*Generalised discrete model*: We now implement an algorithm that can be used to simulate both extensions 1 and 2. In this generalised model, a proliferative agent can place a daughter agent at any vacant target site in 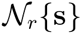, and crowding effects are modeled by *f* (*C*). Therefore, during a potential proliferation event, a randomly selected agent at site s attempts to proliferate with probability *P_p_* per time step of duration *τ*. If the agent is to attempt to proliferate, crowding effects are incorporated by calculating *f* (*Ĉ_s_*). If the potential proliferation event is to succeed, a daughter agent is placed at a randomly selected vacant site in 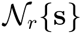.

Figure 2 shows four different schematic illustrations of the generalised proliferation mechanism on a hexagonal lattice. In each illustration, three examples are shown in which sites in 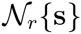 are either: (i) all vacant with *Ĉ*_s_ = 0 (Figure 2(a), (e), (i) and (m)); (ii) half occupied with *Ĉ*_s_ = 0.5 (Figure 2(b), (f), (j) and (n)); or, (iii) fully occupied with *Ĉ*_s_ = 1 (Figure 2(c), (g), (k) and (o)). The first row (Figure 2(a)–(d)) corresponds to the traditional model with *r* = 1 and *f*(*C*) = 1 – *C*. In this case, with *Ĉ*_s_ = 0.5, the probability that a potential proliferation event takes place is *P_p_*/2, and the probability of a particular proliferation event producing a daughter agent at a particular site is *P_p_*/6 (Figure 2(b)). The second row (Figure 2(e)–(h)) corresponds to *r* = 2 and *f*(*C*) = 1 – *C*. In this case, with *Ĉ*_s_ = 0.5, the probability that a potential proliferation event takes place is *P_p_*/2, and the probability of a particular proliferation event producing a daughter agent at a particular site is *P_p_*/18 (Figure 2(f)). Comparing the outcomes in the first and second row of Figure 2 illustrates the first generalisation of the discrete proliferation mechanism as we are simply applying the same proliferation mechanism, with the same crowding function, over a larger template of lattice sites.

The schematic illustrations in the third (Figure 2(i)–(l)) and fourth (Figure 2(m)–(p)) rows of Figure 2 show how the outcomes in the first and second rows can be generalised by choosing different *f* (*C*). For example, with *f*(*C*) = (1 – *C*)^2^, agents are less likely to proliferate than when *f* (*C*) = 1 – *C*. With *r* =1, *Ĉ*_s_ = 0.5 and *f* (*C*) = (1 – *C*)^2^ (Figure 2(j)), the probability that a potential proliferation event takes place is *P_p_*/4, and the probability of a particular proliferation event producing a daughter agent at a particular site is *P_p_*/12, and this is very different to the traditional model (Figure 2(b)).

## III. CONTINUUM DESCRIPTION

While the individual-level details of the generalised discrete proliferation mechanism, highlighted in Figure 2, are very different to the traditional proliferation mechanism, it is not obvious how these differences affect the collective behaviour of a population of agents. To investigate this issue, we will derive the mean field continuum limit description of the discrete models, and then compare the performance of the continuum limit descriptions with averaged data from repeated discrete simulations.

*Traditional model*: We first derive the continuum limit description of the traditional model before considering the more general case. We average the occupancy of site s over many identically prepared realisations to obtain 〈*C*_s_〉 ∈ [0,1] [18] and then develop an approximate discrete conservation statement describing the change in average occupancy of site s from time *t* to time *t* + *τ*,

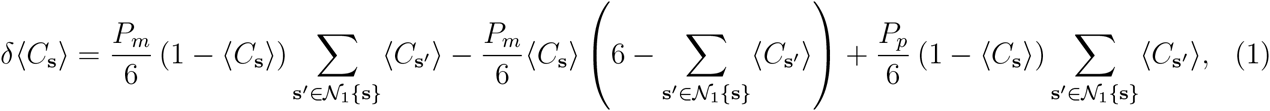

where 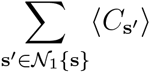 is the sum of the average occupancy of sites in 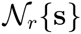. The first and second terms on the right of Equation (1) represent the effects of migration into, and out of, site s, respectively. The third term on the right of Equation (1) represents the effect of proliferation. To arrive at Equation (1), we make the usual mean field assumption that the occupancy of lattice sites is independent as we interpret the products of terms like 〈*C*_s_〉 and 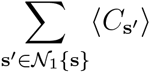 as a net transition probability [17, 18, 29].

We expand each term in Equation (1) as a Taylor series about site s, neglect terms of 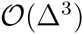, and divide both sides of the resulting expression by *τ*. Identifying 〈*C*_s_〉 with a smooth function, *C*(*x,y,t*), we consider the limit as Δ → 0 and *τ* → 0 jointly, with the ratio *Δ*^2^/*τ* held constant, giving [18]

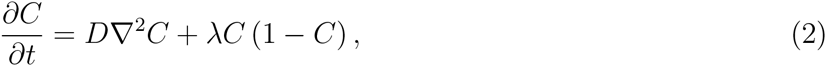

where the diffusivity is 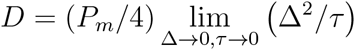, and the proliferation rate is 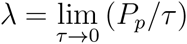. To obtain a well-defined continuum limit we require 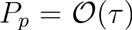 [13, 18]. Equation (2) confirms that the traditional proliferation mechanism is associated with the standard logistic source term. Furthermore, in one dimension, Equation (2) simplifies to the Fisher-Kolmogorov model [30].

*Generalised model:* The migration mechanism in the traditional and generalised models are equivalent, whereas the proliferation mechanism is different. The corresponding approximate discrete conservation statement is

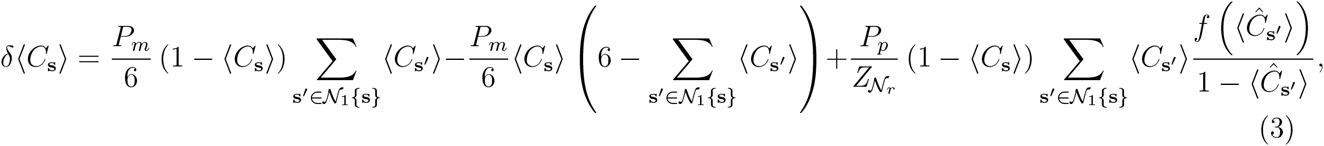

where 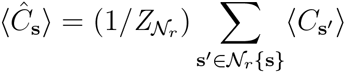. The first two terms on the right of Equation (3) are identical to the corresponding terms in Equation (1). The third term on the right of Equation (3) represents the change in occupancy of site s due to proliferation. The factor 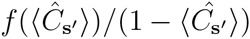 is a measure of the crowding at site s′, in terms of *f* (*C*), relative to the probability that sites in the neighbourhood are vacant. Following the same procedure used previously for the traditional model we obtain,

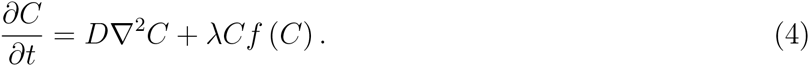

Details of the Taylor series expansions used to derive Equation (4) are given in the Supplementary Material. Comparing Equations (2) and (4), the population-level impact of the change in the proliferation mechanism is to alter the per capita growth rate from the linearly decreasing function of density, λ(1 – *C*), to the more general λ*f* (*C*). For the remainder of this work we set *f* (*C*) = *C*^*α*−1^(1 ‒ *C^β^*)^*γ*^, where *α, β* and *γ* are positive constants [25], but many other choices of *f* (*C*) are possible.

## IV. RESULTS AND DISCUSSION

Our main result, so far, is to describe how to incorporate a generalised proliferation mechanism into a discrete two-dimensional model of cell migration and cell proliferation with crowding effects, and to derive the mean field continuum limit description. However, at this stage, it is unclear how well the continuum model will predict averaged data from repeated stochastic simulations of the discrete model. To explore this issue we now apply the discrete and continuum models to mimic both a cell proliferation assay and a scratch assay (Figure 1). We systematically vary *r* and *f* (*C*) to explore how these choices affect the performance of the continuum description.

A key parameter in the discrete model is *P_p_/P_m_*, which is the relative frequency of proliferation events to motility events for an isolated agent [18]. This ratio can be estimated from experimentally observable quantities including the doubling time 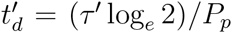, the cell diffusivity *D*′, and the average cell diameter, Δ′. For typical values: 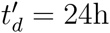 [18]; D′ = 1000 *μ*m^2^/h [5]; and, Δ′ = 24*μ*m [6], we have *P_p_/P_m_* ≈ 0.001. Therefore, all simulations and analysis in the main document correspond to Δ = *τ* = *P_m_* = 1 and *P_p_* = 0.001. These non-dimensional simulations can be used to model a population of cells with an arbitrary dimensional cell diameter and an arbitrary doubling time by re-scaling Δ and *τ* with appropriate choices of Δ′ and *τ*′, respectively [18]. To ensure the conclusions drawn from these simulations are applicable to a wide range of cell lines, we repeat all simulations and analysis for a higher proliferation rate, Δ = *τ* = *P_m_* = 1 and *P_p_* = 0.05 (Supplementary Material).

*Cell proliferation assay*. We consider a suite of simulations of a cell proliferation assay based on the geometry of the images in Figure 1(a)–(c). We use a lattice of size I x J to accommodate a typical population of cells (Δ′ = 24*μ*m [6]). Since the images in Figure 1(a)–(c) show a fixed field of view that is much smaller than the spatial extent of the real experiment, and the cells in the experiment are distributed uniformly, we apply zero net flux boundary conditions along all boundaries [31]. Simulations are initiated by randomly populating each lattice site with a constant probability of 5%, so that there are, on average, no spatial gradients in agent density across the lattice. Snapshots from the model, across a range of choices of *f*(*C*), are given in Figure 3.

**Figure 3.**
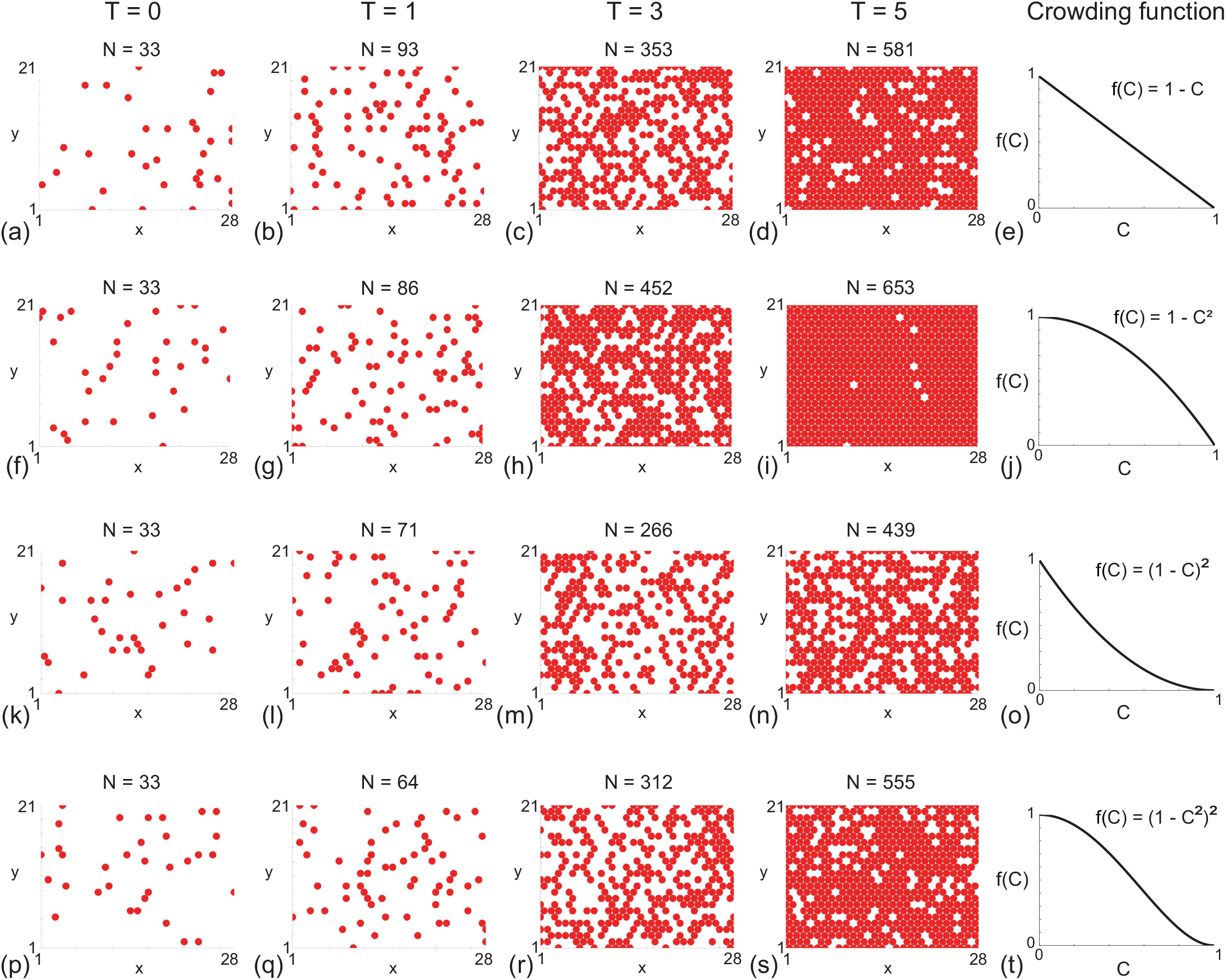
Snapshots of simulations for a suite of cell proliferation assays. In each row the distributions of agents at time λ*t* = *T* = 0,1,3,5, as indicated, are shown along with the corresponding *f(C)*. Each simulation is initiated by randomly populating a lattice of size *I* = 28 and *J* = 24, so that each site is occupied with probability 5%. All simulations correspond to Δ = *τ* = *P_m_* = 1, *P_p_* = 0.001 and *r* = 4.

To mimic the way that cell proliferation assays are reported [4] (Figure 1(d)), we calculate the time evolution of the total number of agents on the lattice, which, when divided by the total number of lattice sites, gives the agent density per unit area [18]

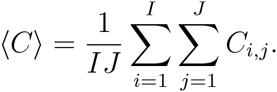

We further average these results over many identically prepared simulations so that we report relatively smooth data where stochastic fluctuations in the agent density are negligible. To compare averaged simulation data with the solution of the continuum model, we note that the absence of spatial gradients means that, on average, ∇^2^*C* = 0. Therefore, instead of dealing with a PDE for *C*(*x,y,t*), Equation (4) simplifies to the ordinary differential equation (ODE) for *C*(*t*) [4]

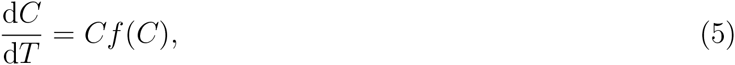

where we have written *T* = *t*λ. This re-scaling of the time variable allows us to more easily compare results in the main paper for a standard proliferation rate (*P_p_/P_m_* = 0.001) with additional results for faster proliferation (*P_p_/P_m_* = 0.05) (Supplementary Material). We solve Equation (5) numerically using a backward Euler approximation with a constant time step, *δt*, and Picard linearisation with convergence tolerance, *ϵ*.

Results in Figure 4(a)–(b) show that when *f*(*C*) is linear, the discrete and continuum density profiles are indistinguishable at this scale for *r* = 1, 2, 3 and 4. Results in Figure 4(c) quantify the discrepancy between the solution of the continuum model and averaged discrete density data using *E* = 〈*C*〉 − *C*, where 〈*C*〉 is the average density per unit area from the discrete simulations and *C* is the solution of Equation (5). In summary, the evolution of *E* (Figure 4(c)) shows that the error is extremely small, with no discernible trends for the different choices of *r*. Additional results in Figures 4(d)–(l) show similar comparisons for a range of nonlinear *f*(*C*) and several choices of *r*. The averaged discrete data and the solution of the corresponding continuum model show that, broadly speaking, the continuum model provides a good prediction of the averaged discrete results but there are some differences for different choices of *r* when *f*(*C*) is nonlinear. When we compare the evolution of the density data between Figures 4(b), (e), (h) and (k), we see that the choice of *f*(*C*) impacts the evolution of the density profile. Similar comparisons to those in Figure 4 are made in the Supplementary Material document for a higher initial condition. In addition to these two different choices of initial condition, we also performed comparisons for a range of other initial conditions (not shown) and these results confirm that the trends we observe are relevant regardless of the initial condition.

**Figure 4.**
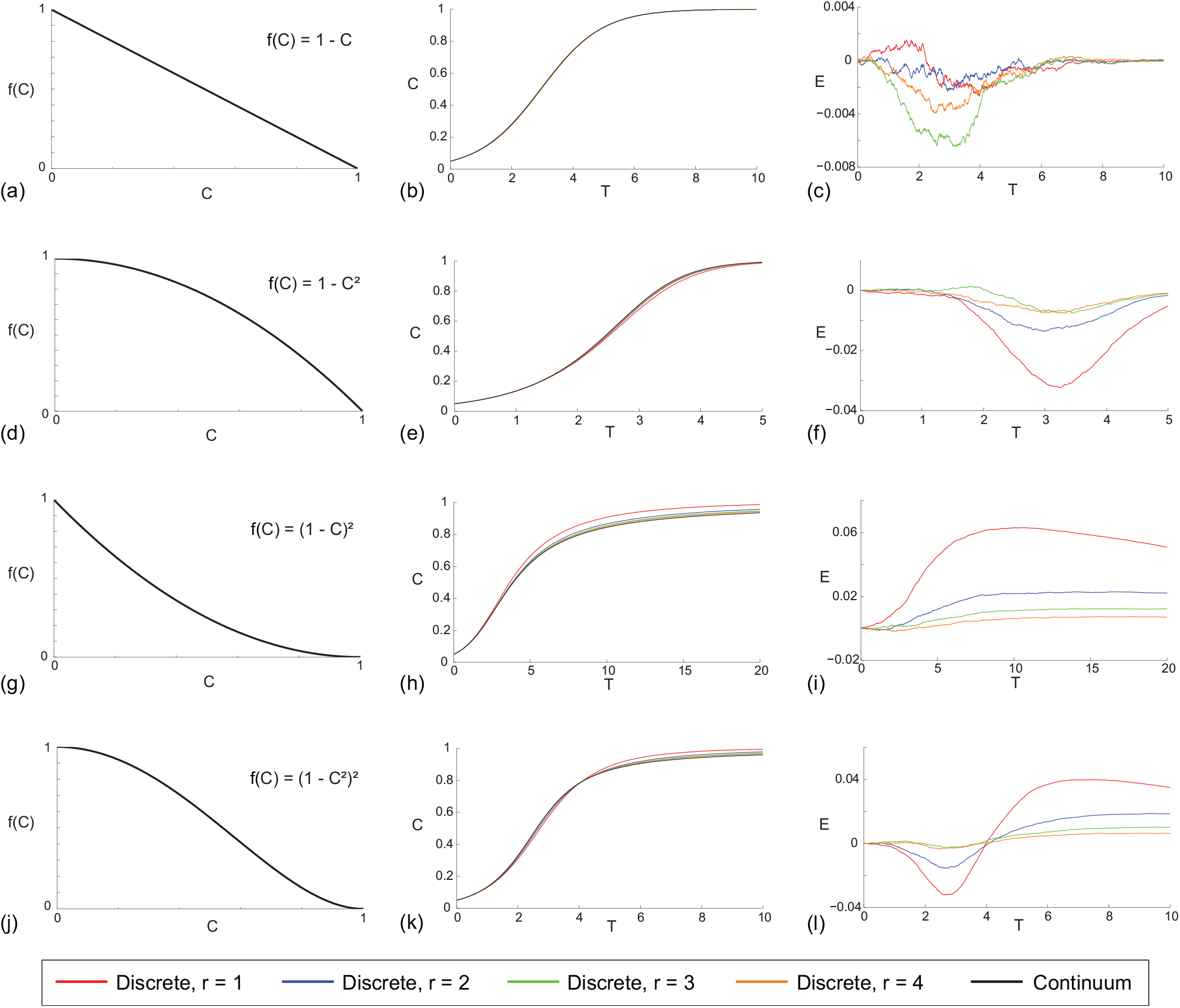
Comparison of averaged simulation data and the solution of the corresponding continuum model for a cell proliferation assay with: *f*(*C*) = 1 − *C*, as shown in (a)-(c); *f*(*C*) = 1 − *C*^2^, as shown in (d)-(f); *f*(*C*) = (1 − *C*)^2^, as shown in (g)-(i); and *f*(*C*) = (1 − *C*^2^)^2^, as shown in (j)-(l). Results in (b), (e), (h) and (k) compare averaged simulation data and the solution of the corresponding continuum model for a range of 1 ≤ *r* ≤ 4 where the initial condition corresponds to 5% of sites being randomly occupied. All simulations are performed on a lattice with *I* = 28 and *J* = 24, and results are averaged across 300 identically prepared realisations of the discrete model. Profiles in (c), (f), (i) and (l) quantify the discrepancy between the solution of the continuum model and the average simulation data. All simulation results correspond to Δ = *τ* = *P_m_* = 1 and *P_p_* = 0.001, and the numerical solution of the continuum model is obtained with δ*t* = 1 × 10^−3^ and *ϵ* =1 x 10^−5^. In all cases *T* = λ*t*

Although the match between the average discrete data and the solution of the corresponding continuum model in Figure 4 is very good, there are some trends that are not obvious without making these comparisons explicit. For example, results in Figure 4(e)–(f), (h)-(i) and (k)-(l) indicate that the performance of the continuum model is slightly poorer when *f*(*C*) is nonlinear compared to the results in Figure 4(b)–(c) where *f*(*C*) is linear. However, for all choices of *f*(*C*), the performance of the continuum model improves as *r* increases. For example, all results with *r* = 4 lead to an excellent match regardless of *f*(*C*). Therefore, these results indicate that estimating r from experimental time lapse images, such as those reported by Druckenbrod and Epstein [28], will be important if we need to decide whether the continuum approximation is sufficient and r is sufficiently large, or whether we need to use more computationally demanding repeated simulations of the discrete model, when *r* is sufficiently small.

*Scratch assay*. We also consider a suite of simulations of a scratch assay based on the geometry of the images in Figure 1(e)–(g). Since these images show a fixed field of view that is much smaller than the spatial extent of the real experiment [6], we apply zero net flux boundary conditions along all boundaries of the lattice [31]. We model the scratch assays on a lattice of size *I × J* that is chosen to accommodate a typical population of cells (Δ′ = 24*μ*m [6]). To model the initial condition, we randomly populate all lattice sites with an equal probability of 30%. This initial density is smaller than the maximum carrying capacity density, and this is consistent with previous experimental procedures [6]. While we present results for an initial density of 30%, our stochastic simulation tools and continuum limit description are general, and can deal with any initial density. To simulate the scratch, we remove all agents from a vertical region, with a width of 23 cell diameters (Figure 1(d)). Snapshots from the discrete model, for a range of *f*(*C*) are shown in Figure 5.

**Figure 5.**
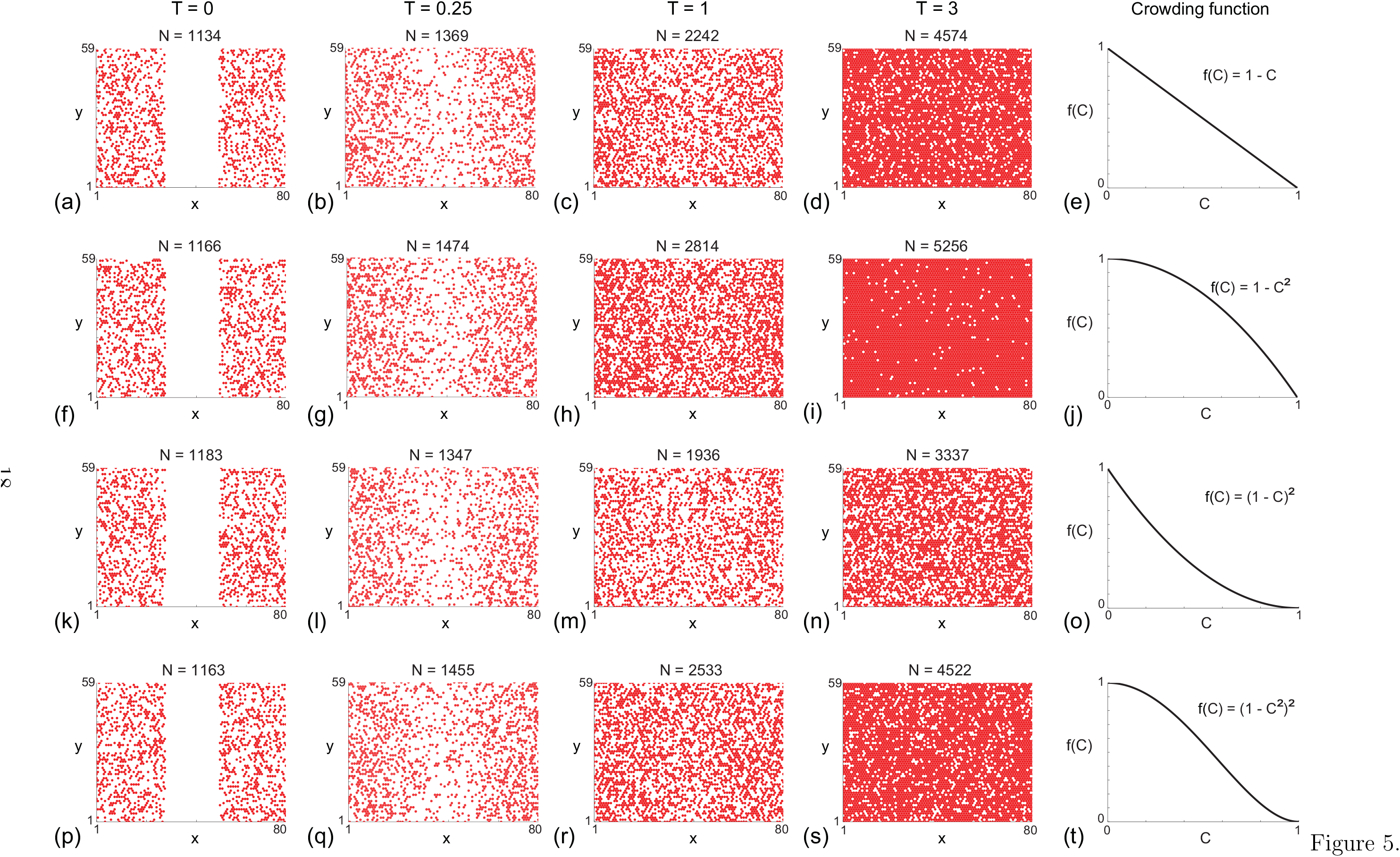
Snapshots of simulations for a suite of scratch assays. In each row the distributions of agents at time λ*t* = *T* = 0, 0.25,1, 3 are shown along with the corresponding *f(C)*. Each simulation is initiated by randomly populating a lattice, corresponding to lattice of size *I* = 80 and *J* = 68, so that each site is occupied with probability 30%. A scratch of 23 lattice sites wide is made at *T* = 0. All simulations correspond to Δ = *r* = *P_m_* = 1, *P_p_* = 0.001 and r = 4.

Since the initial condition is uniform in the vertical direction, we average the agent population density along each vertical column of lattice sites to obtain

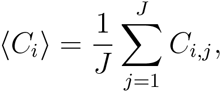

which is then further averaged over many identically prepared discrete simulations to reduce fluctuations. This procedure allows us to plot the time evolution of the average agent density as a function of the horizontal coordinate, as shown in Figure 1(h) [6, 31]. Typical results from the discrete model are shown in Figure 6(a)–(c), with the standard linear *f*(*C*), for *r* =1 and *r* = 4, respectively. As time increases, we see the effects of combined agent motility and agent proliferation as the agent density profiles spreads into the initially-vacant region. The effects of proliferation can also be observed as the density profile increases with time towards confluence.

**Figure 6.**
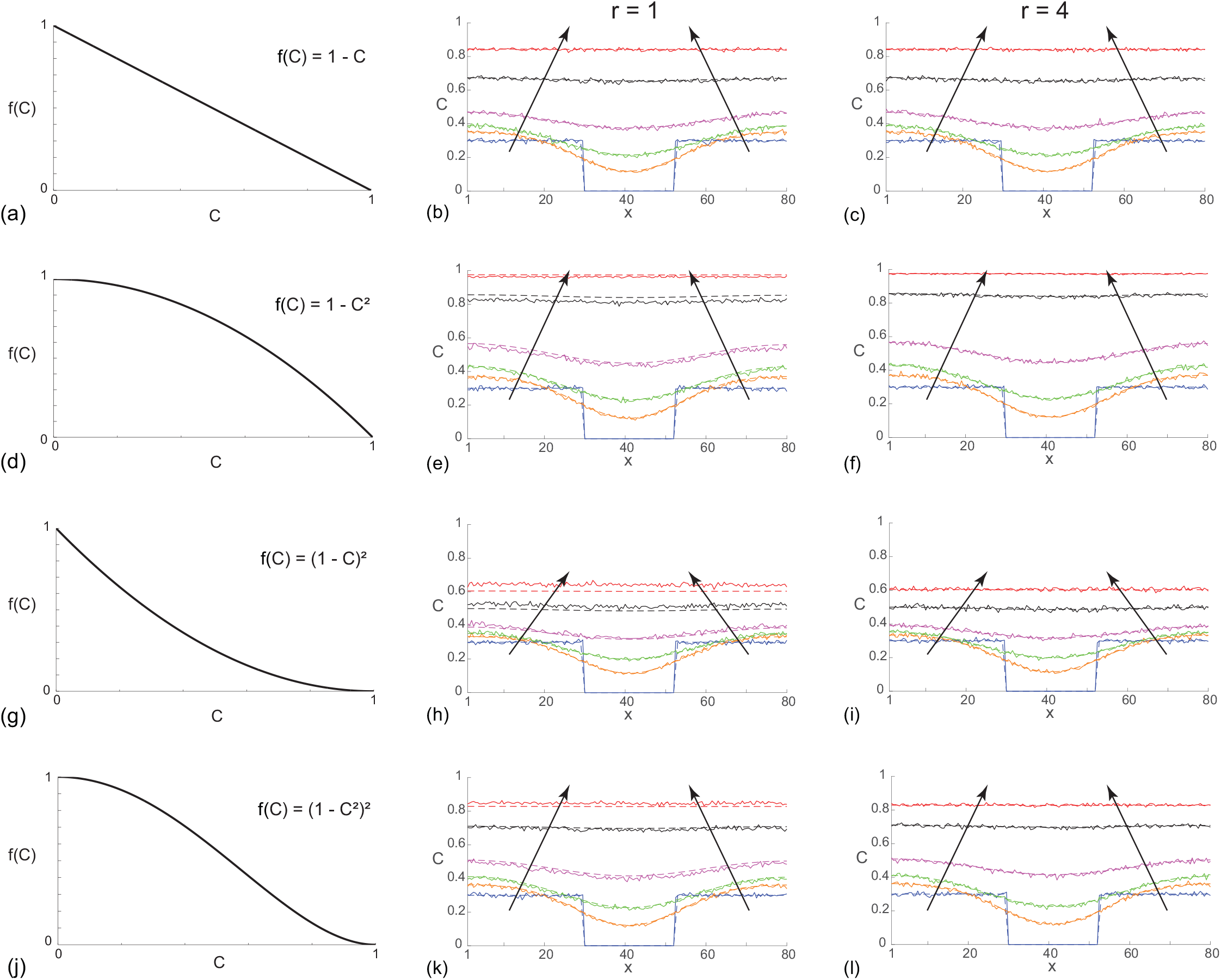
Comparison of averaged simulation data and the solution of the corresponding continuum model for a scratch assay with: *f*(*C*) = 1 − *C*, as shown in (a)-(c); *f(C)* = 1 − *C*^2^, as shown in (d)-(f); *f(C)* = (*f* − *C*)^2^, as shown in (g)-(i); and *f*(*C*) = (1 − *C*^2^)^2^, as shown in (j)-(l). Results in (b), (e), (h) and (k) compare averaged simulation data (solid lines) and the solution of the corresponding continuum model (dashed lines) for *r* = *f*. In each subfigure, agent density profiles are given at λ*t* = *T* = 0, 0.25, 0.5, 1, 2, 3, and the direction of increasing *t* is shown with the arrows. Results in (c), (f), (i) and (1) show an equivalent comparison except here we have *r* = 4. All simulation results are averaged across 100 identically prepared realisations of the discrete model, with Δ = *r* = *P_m_* = 1 and *P_p_* = 0.001, on a lattice of size *I* = 80 and *J* = 68. The numerical solution of the continuum model is with *δx* = 0.25, *δt* = 0.1 and *ϵ* = 1 × 10^−5^.

To explore how well the continuum description matches this vertically-averaged discrete density data, we note that since the initial condition is independent of the vertical location (Figure 1(d)), we can average Equation (4) in the vertical direction [6, 31] to give,

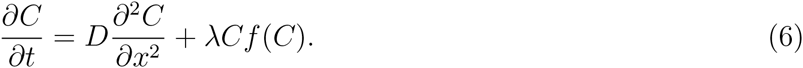

To solve Equation (6), we apply no flux boundary condition at both boundaries, and the initial condition is given by *C*(*x*, 0) ≡ 0 within the initially-vacant region, and *C*(*x*, 0) ≡ 0.30 outside of this region. We solve Equation (6) numerically using a central difference approximation with uniform spacing *δx*, and the temporal derivative is approximated using a backward Euler method with a uniform time step, *δt*. The resulting system of nonlinear algebraic equations is solved using Picard iteration with convergence tolerance, e. The numerical solution of Equation (6) is superimposed on the averaged discrete density profiles in Figure 6(b)–(c) for *f*(*C*) = 1 − *C*, with *r* =1 and *r* = 4, respectively. As with the cell proliferation assay results in Figure 4, the quality of the continuum-discrete match for the scratch assay with *f*(*C*) = 1 − *C* is excellent for all *r* considered. Data in Figure 6(d)–(l) show similar comparisons for simulations of scratch assays for a range of nonlinear *f*(*C*). Comparing the evolution of *C*(*x,T*) in Figure 6 shows how the choice of *f*(*C*) influences the evolution of the scratch assay. Again, as for the cell proliferation assays, overall we observe a good quality of match between the solution of the continuum model and the averaged agent density profiles across all choices of *f*(*C*) and r for the scratch assays. However, for nonlinear *f*(*C*), we observe some small discrepancies, and these discrepancies are most pronounced when *r* =1. In contrast, the quality of the continuum-discrete match is excellent for larger r across all choices of *f*(*C*) that we consider. Further comparisons of the performance of the continuum PDE description for intermediate values of *r* = 2 and *r* = 3 are given in the Supplementary Material document.

## V. Conclusions

Most continuum [3, 7–10] and discrete [5, 17, 18] models of collective cell spreading are associated with a logistic growth source term to model cell proliferation. However, certain *in vivo* [23, 24] and *in vitro* [6] evidence suggests that cells do not always proliferate logistically. Motivated by these observations, we extend the traditional exclusion process based discrete model of cell proliferation since the continuum limit description of this traditional model always leads to logistic growth. Our new, generalised discrete model encompasses two extensions. The first extension involves choosing a variably sized proliferation template so that the target site is chosen from a set of sites contained within *r* ≥ 1 concentric rings about the mother agent. The second extension involves a measure of the crowdedness of the proliferative agent, *Ĉ*_s_ ∈ [0,1]. A *crowding function*, *f*(*C*), which incorporates information from a group of neighbouring sites is used to quantify the influence of crowding. Analysing the mean field continuum limit of the generalised model shows that the usual logistic source term, λ*C*(1 − *C*), is generalised to λ*Cf*(*C*). There are several interesting consequences of this generalised mean field PDE, namely: (i) the traditional logistic source term corresponds to linear *f*(*C*); (ii) the size of the template, *r*, does not appear in the continuum limit description; and, (iii) without making explicit comparisons, it is unclear how different choices of *f*(*C*) and *r* affect the accuracy of the continuum limit description.

To provide insight into how different choices of *f*(*C*) and *r* affect the accuracy of the continuum limit description, we generate averaged discrete data from the generalised random walk model for both a cell proliferation and a scratch assay for a typical cell line with *P_p_/P_m_* = 0.001 [5]. Averaged simulation data are generated for a range of choices of *f*(*C*) and *r* and we find that, overall, the continuum description provides a good prediction of the average behaviour of the stochastic simulations. While there is a modest discrepancy in the continuum-discrete match for nonlinear *f*(*C*), we find that the quality of the match improves as *r* increases. Therefore, to make a distinction between the need for using repeated stochastic simulations or simply working with the continuum description, we suggest that experimental time lapse images [28] ought to be used to provide an estimate of *r*. To ensure that our conclusions are broadly applicable across a wide range of cell lines, all continuum-discrete comparisons are repeated for a cell line with a particularly fast proliferation rate *P_p_/P_m_* = 0.05 (Supplementary Material), and we find that the same conclusions apply.

There are two features of this study that could warrant further investigation. First, since the focus of the work is to investigate the role of the proliferation mechanism, all simulations and analysis invoke the most fundamental unbiased nearest neighbour exclusion motility mechanism. This mechanism provides a good approximation of the collective motility of mesenchymal cell lines that are largely unaffected by cell-to-cell adhesion [5, 7]. However, if dealing with an epithelial cell line, it would be more reasonable to invoke a motility mechanism that incorporates cell-to-cell adhesion [29, 32]. Under these conditions it would be interesting to extend the present analysis to investigate the performance of the continuum limit description of the generalised proliferation mechanism with an adhesive motility mechanism [29, 32]. Second, all discrete simulations in this work are lattice-based. While many previous studies have used lattice-based models to successfully mimic and predict a range of two-dimensional *in vitro* assays [7, 20, 31, 32], all lattice-based models make certain implicit assumptions. For example, all lattice-based exclusion process models effectively assume that agents are a fixed size [15–20], which is clearly an approximation because it is well known that cell proliferation involves gradual changes in cell volume. An alternative way to simulate collective cell migration experiments would be to use a lattice-free framework [33, 34]. One of the advantages of using a lattice-free framework is that the model can be adapted to allow for dynamic cell size changes during a proliferation event. However, the main limitation of using a lattice-free method to mimic a real experiment is that the computation time is proportional to *N*^2^, where *N* is the number of agents in the simulation. This can be prohibitive if we wish to mimic a real *in vitro* experiment with many tens of thousands, or hundreds of thousands cells [32]. In contrast, lattice-based methods are more convenient since the computation time is proportional to *N*.

## Acknowledgments

This work is supported by the Australian Research Council (DP140100249, FT130100148). We thank Parvathi Haridas and Esha Shah for providing the experimental images in Figure 1, and we appreciate the helpful comments from the three referees.

